# Sex specific role of RNA-binding protein, AUF1, on prolonged hypersensitivity after repetitive ischemia with reperfusion injury

**DOI:** 10.1101/2023.06.08.544080

**Authors:** Meranda M. Quijas, Luis F. Queme, Alex A. Weyler, Ally Butterfield, Diya P. Joshi, Irati Mitxelena-Balerdi, Michael P. Jankowski

**Author notes:** ^†^Current Affiliation: Department of Biomedical Sciences, University of New England. **Corresponding author:** Michael P. Jankowski 3333 Burnet Avenue, MLC6016 Cincinnati, Ohio 45229-3026.

## Abstract

Repetitive ischemia with reperfusion (I/R) injury is a common cause of myalgia. I/R injuries occur in many conditions that differentially affect males and females including complex regional pain syndrome and fibromyalgia. Our preclinical studies have indicated that primary afferent sensitization and behavioral hypersensitivity due to I/R may be due to sex specific gene expression in the DRGs and distinct upregulation of growth factors and cytokines in the affected muscles. In order to determine how these unique gene expression programs may be established in a sex dependent manner in a model that more closely mimics clinical scenarios, we utilized a newly developed prolonged ischemic myalgia model in mice whereby animals experience repeated I/R injuries to the forelimb and compared behavioral results to unbiased and targeted screening strategies in male and female DRGs. Several distinct proteins were found to be differentially expressed in male and female DRGs, including AU-rich element RNA binding protein (AUF1), which is known to regulate gene expression. Nerve specific siRNA-mediated knockdown of AUF1 inhibited prolonged hypersensitivity in females only, while overexpression of AUF1 in male DRG neurons increased some pain-like responses. Further, AUF1 knockdown was able to specifically inhibit repeated I/R induced gene expression in females but not males. Data suggests that RNA binding proteins like AUF1 may underlie the sex specific effects on DRG gene expression that modulate behavioral hypersensitivity after repeated I/R injury. This study may aid in finding distinct receptor differences related to the evolution of acute to chronic ischemic muscle pain development between sexes.

## Introduction

Musculoskeletal pain is a major health concern and is experienced more than any other type of pain worldwide [3,16,21,35,37,48]. Chronic musculoskeletal pain (myalgia) specifically, defined as pain persisting beyond normal healing time, is a prevalent burden causing years of disability [6]. Pain is often widespread throughout the body, making strategies to treat the underlying issue difficult. Myalgia can be due to numerous underlying causes: overuse or strain of muscle, traumatic injury, or disease. A major factor of disease-based myalgia is ischemia with reperfusion (I/R) injury. Repeated I/R injury occurs in many disorders such as peripheral vascular disease, sickle cell anemia, or even fibromyalgia and complex regional pain syndrome [1,12,17,25,51,65].

Diseases like these are often reported more in the female population, where pain tolerance is also lower [11,15,19,38,48,68,69]. The mechanisms of sexually dimorphic pain hypersensitivity have been linked in part to the immune system [34,52] with far less known about sex differences in the peripheral nervous system. At the cellular level, there is emerging evidence that different male and female mechanisms may govern pain responses [34,42,51].

Although animals of both sexes show similar pain-like behaviors following I/R injury, analysis of the dorsal root ganglia (DRGs) showed that I/R evoked an increase in acid-sensing ion channels (ASIC) 1 and 3 in males, while females showed an upregulation of transient receptor potential vanilloid type 1 (TRPV1) as well as TRP Melastatin 8 (TRPM8) [55]. This suggests that distinct molecular pathways may underlie pain-like behaviors after I/R between males and females.

One possible factor that may affect the sex-specific mechanisms within DRG neurons leading to prolonged hypersensitivity could involve RNA binding proteins which modulate gene expression [7,26]. One such protein is AU-rich element RNA binding protein factor 1 (AUF1, aka, HNRNPD) which has been shown to regulate cytokine-related mRNAs and can have either stabilizing or degrading effects, depending on the isoform [56,70]. AUF1 binds transcripts encoding cytokine receptors and plays a role in inflammatory responses [66], which have been suggested to be important regulators of female specific responses to I/R [55]. Here we wanted to investigate the role of sensory neuron AUF1 in a model of prolonged hypersensitivity after repeated I/R injury. Highlighting the potential sex specific mechanisms leading to pain under these conditions could be beneficial for future preclinical and clinical studies and lead to novel sex specific therapies for ischemic myalgia.

## Materials and Methods

### Animals

Adult male and female Swiss Webster mice (Charles River) 3-6 weeks of age were used throughout all experiments. Mice were housed in a barrier facility maintained on a 14:10-h light-dark cycle with a temperature-controlled environment and given food/water ad libitum. All procedures were approved by the Cincinnati Children’s Hospital Institutional Animal Care and Use Committee and adhered to NIH Standards of Animal Care and Use under the Association for Assessment and Accreditation of Laboratory Animal Care International (AAALAC)-approved practices.

### Ischemia with Reperfusion (I/R) Injury

Mice were anesthetized with 2-3% isoflurane for surgical procedures. For I/R injuries, the brachial artery in the right forelimb was exposed and the artery was ligated using a 7.0 silk suture. Incisions were closed with a 6.0 silk suture after the occlusion. After 6 hours, animals were re-anesthetized for removal of the arterial occlusion suture. A second I/R injury was then performed 7 days after the first. Sham mice were also utilized by performing an incision and loosening of the vessel on each surgery day without performing the occlusion.

### Interventions

To determine the most effective siRNA used for *in vivo* experiments, knockdown efficiency was tested via neuro-2A cells *in vitro*. In a 12 well plate, approximately 10^4^ cells/well were seeded and incubated overnight (18-24hours). TransIT-TKO (Mirus) transfection reagent was prepared adding 2.5μL to 100µl of media without serum. Next, 10 µM siRNA stock solution from one of four different targeting duplexes or a non-targeting control was added to the transfection reagent mixture to create a 10nM final concentration per well. This was incubated at room temperature for approximately 20 minutes. Then each of the four siRNA duplexes (or control duplex) were distributed dropwise into separate wells containing cells in complete growth medium (Eagle’s Minimum Essential Medium, 10% Fetal Bovine Serum, and 1% Penicillin-streptomycin). Cells were incubated for approximately 24 hours and harvested for RT-PCR analysis to determine knockdown efficiency. The AUF1 siRNA found to have the greatest targeting efficiency *in vitro* **(Supplementary Fig. 1)** was modified for binding to the lipophilic peptide, Penetratin-1 similar to our previous reports [e.g.49,50,54]. To knockdown AUF1 mRNA *in vivo* in mice with dual I/R injury, animals were injected with 0.1-0.2 µL of Penetratin-linked AUF1 (PenAUF1) targeting siRNAs or non-targeting controls (PenCon) into the median and ulnar nerves [50,54] on days 4 and 5 after the initial I/R using a quartz microelectrode connected to a picospritzer. Wounds were then closed with 6.0 silk sutures.

To overexpress AUF1 in select peripheral nerves *in vivo*, a similar injection procedure was followed except adeno-associated viruses, serotype 9 (AAV9), containing an AUF1 overexpression construct were injected into the median and ulnar nerves 3-4 weeks prior to the first I/R injury. AAVs with AUF1 the overexpression construct (AAV9-CAG-m-HNRNPD-IRES-eGFP, SKU: AAV-261522) or controls (AAV9-CAG-EGFP, Cat No: 7076) were purchased from Vector Biolabs and injected into the median and ulnar nerves in 1uL volumes as described above. After injections, incision sites were closed using 6.0 silk sutures.

### Behavioral Assays

Mice were tested for baseline behavior one day before injury as described previously [50,54]. Mice were analyzed at baseline prior to the initial I/R and then again at day 1 and 6 post I/R. After the second I/R, animals were again tested at days 1, 3, 6 and 8 post second injury. Each day, two behavioral tasks were performed: assessment of spontaneous paw guarding (non-evoked) and analysis of mechanical withdrawal thresholds to fore paw muscle squeezing (evoked). Mice were first placed individually in raised chambers with a mesh bottom and allowed to habituate for at least 15 minutes. Fore paw guarding was conducted by observing the mice for one hour, at five-minute intervals, assigned a score from 0-2 (0-mouse using all paws without reservation, 1-mouse isn’t bearing all weight on paws equally, 2-mouse holds paw completely off the mesh) after observation for 1 min. The average of the 12 trials was determined each day and then averaged across all mice within an experimental group. Mechanical hypersensitivity was performed on both forepaws using a set of digital calibrated forceps with a blunt probe. Mice were held and forepaws squeezed with increasing force until a withdrawal response was noted. The force at which the animal withdrew was recorded. The cut-off intensity was set at 300g. Withdrawal thresholds were determined as a set of three trials, performed three times with a five-minute interval. The average of the three trials were used for analysis. In addition to this, fore paw grip strength was also analyzed in separate cohorts using a grip strength meter (BioSeb, France). Animals were held by the tail above a metal grid and allowed to grip the grid with only their forepaws. They were then pulled back horizontally until they could no longer retain their grip. The grip strength was measured in grams for three rounds of three trails with 5 minutes in between. The average of all nine trials was taken per testing day and corrected for weight differences for analysis. The experimenter was blinded to the treatment groups during all behavioral assays.

### Proteomics Analysis

DRGs (C7, C8, & T1) were collected from both male and female Swiss Webster mice and samples were run through comparative profiling to identify protein changes via SWATH nanoLC-MS/MS using runs from each sample to generate quantitative comparisons. The frozen tissue was placed in an Eppendorf homogenizer (Kontes749520-0000) and then 25 µl of homogenization buffer (10mM Tris with Roche complete protease inhibitor) was added and the sample homogenized for several minutes before an additional 10µl of homogenization buffer was added again and re-homogenized. Next, 25 µl of 2X Invitrogen sample buffer without dye was added and the sample was sonicated in a water bath for 1 minute followed by cooling on ice for 15 min for two cycles. The samples were then centrifuged at 1000xg for 10 minutes and the supernatant was placed in a new tube.

Next the Pierce 660nm Protein assay was performed to determine the protein concentration using a BSA as a standard. 20 µg of each sample in 40 µl of Laemmli buffer were run 1.5 cm into a 1D, 1.5 mm 4-12% BT gel using MOPS running buffer. Pre-stained protein markers were used in surrounding lanes and the regions between the markers and the dye front were excised for trypsin digestion following a standard gel protocol. The resulting peptides were extracted, dried and prepped for mass spectrometry. 2.5 µg of each sample was run on the nanoLX-MS/MS in DDA mode and the combined DDA runs were searched using Protein Pilot to create a protein spectral library. From this, 1702 proteins were identified with a false discovery rate (FDR) of less than 1% at peptide and protein levels.

A matched SWATH-MS method in DIA mode of the samples was used to collect quantitative data for each of the samples for comparative profiling (8 runs) and the total ion chromatograms were generated. SVVATH-D data analysis workflow was used to validate the data set and detect any significant quantitative changes. From this, 1175 proteins were identified and quantified with 99% confidence with FDR of less than 1%. A heat map of the 30 proteins with significant changes between the groups was generated (p<0.05) as well as a volcano plot of significant proteins (p value<0.05 and >1.25 fold change) to show upregulated and down regulated proteins.

### Western Blotting

Forepaw muscle was dissected from anesthetized mice and flash frozen in liquid nitrogen followed by storage at -80°C. After homogenization in protein lysis buffer containing 1% SDS, 10mM Tris-HCL (pH 7.4) and protease inhibitors (1µg/mL pepstatin, leupeptin, aprotinin, 1 mM sodium orthovanadate, and 100 µg/ml phenylmethylsulfonyl fluoride; Sigma-Aldrich). 20µg samples were boiled in gel loading buffer containing β-mercaptoethanol and sodium lauryl sulfate as a reducing agent and loaded onto AnyKD precast polyacrylamide gels (Bio-Rad 4569033) for protein separation. Proteins were transferred to polyvinylidene difluoride membranes (PVDF; Merck Millipore Ltd.) at 35 V overnight in a cold room at 4°C. The following day, the membrane was washed, blocked with buffer (Odyssey; LiCor 927-40000) diluted in PBS (1:4) and incubated in 2x PBS with 0.2% Tween and blocking buffer (1:1) with primary antibodies rabbit anti-AUF1 (Abcam, 1:5000), rabbit anti-phospho-AUF1 (Abcam, 1:2000), rabbit anti-HuR (Abcam,1:5000), rabbit anti-phospho-HuR (Abcam, 1:200), mouse anti-IL1β (R&D Systems, 1:500), rabbit anti-GDNF (Alomone, 1:500), and chicken anti-GAPDH (Abcam, 1:2000). After incubation over-night, the membranes were washed and incubated in 2x PBS with 0.2% Tween and 0.01% SDS and blocking buffer (1:4) with appropriate infrared-conjugated secondary antibodies (LiCor IRDye 680RD Donkey anti-Chicken 926-68075 Lot no. C80717-13, IRDye 800CW Donkey anti-Rabbit 926-32213 Lot No. c80929-05, Goat anti-mouse IRDye 800CW Lot No C31021-01). Membranes were visualized on LiCor Odyssey CLx protein imaging system. Exposure times were kept consistent between runs and gain was set to 1.0. Band intensity was quantified using ImageJ software (NIH) via previous procedures [52]. The immunoreactive bands were then analyzed via densitometry and then quantified through ImageJ software. The optical density was normalized to GAPDH and reported as protein quantification.

### Immunohistochemistry

To extract tissue, mice were first anesthetized via an intramuscular injection of Ketamine/Xylazine Solution (9 mg/mL Ketamine + 0.9 mg/mL Xylazine). For DRG extraction, post perfusion with ice cold 0.9% saline solution, a laminectomy was performed to expose the spinal cord and then the C7, C8, and T1 DRGs were isolated. For muscle extraction, skin and digits were removed from the right forepaw and the muscle was isolated and placed in disposable vinyl specimen molds with optimal cutting temperature (O.C.T.) compound.

After tissue dissection, DRGs were immersion fixed in 4% paraformaldehyde for 30 min prior to placement in specimen molds with O.C.T compound while muscle sections were flash frozen in liquid nitrogen. Sections were cut using a cryostat at 7 microns for DRG, 12 microns for muscle tissue. Slides were mounted with subzero freezing medium (Mercedes Scientific) and allowed to dry on a slide incubator for at least 5 minutes. Tissue was fixed on the slides with 4% PFA in 0.01 M PSB for 15 min (2-5 minutes for DRGs). Slides were rinsed in 0.1 M PBS and then underwent antigen retrieval, for 23 minutes in citrate buffer, at 73°C. Slides were then blocked in 0.01 M PBS with 4% donkey serum, 4% horse serum, 1% bovine serum albumin, and 0.01M PBS with 0.1% Tween-20 for 1 hour. Then the slides were incubated in the above blocking buffer overnight at room temperature with gentle rocking with the primary antibodies as indicated guinea pig anti-ASIC3 (1:2000, Millipore), rabbit anti-TRPV1 (1:2000, Alomone), mouse anti-IL1β (1:400, R&D Systems), and rabbit anti-GDNF (1:100, Alomone). Next, slides were washed in 0.01M PBS and incubated in the same blocking buffer but with appropriate secondary antibodies (647 donkey anti-guinea pig, 1:400, 488 donkey anti-mouse and 594 donkey-anti rabbit, Jackson Immunoresearch). The slides were then rinsed in 0.01M PBS and cover slipped using Fluro media with DAPI to stain nuclei. Labeling of cells was identified and characterized on a Nikon A1 inverted confocal microscope with sequential scanning. Images were then compiled and prepared for publishing via NIS-elements imaging software. For quantification, non-sequential sections of samples were counted to prevent any overlapping cells from being quantified more than once. Cells that were positive for one or more antibodies were counted separately and each marker was quantified for each section. Counts were performed on 6-8 sections per single animal and the average counts per condition were used for analysis (n=3-4).

### Real-time RT-PCR

RNA was isolated from C7, C8, T1 DRGs using Qiagen RNeasy Mini kits according to the manufactures’ directions. 500ng of total RNA was reverse transcribed into cDNA using Superscript II and 2 ng of cDNA per well was used with SYBR-Green real-time PCR reactions on a StepOne real-time PCR System (Applied Biosystems). Ct values were obtained and analyzed by the ΔΔCt method after normalization to GAPDH. Expression differences are determined from the normalized ΔΔCt values and standard error of the difference in means. Fold change between conditions is determined and values were converted to a percent change where 2-fold = 100% change [50].

### Experimental design and statistical analysis

Data were analyzed using GraphPad Prism 9.5.0. Critical significance was set to α < 0.5. All data were checked for normal distribution and equal variance using the Shapiro-Wilk normality tests and then parametric or nonparametric tests were used accordingly. All behavioral tests are specifically indicated in the figure legends. Behavioral analyses were analyzed via a two-way repeated measures ANOVA with Holm-Sidak and/or Tukey’s post hoc tests. Analysis of protein quantification from western blots were measured via one-way ANOVA or corresponding non-parametric test across groups. RT-PCR was analyzed via one-way ANOVA with Holm-Sidak post hoc test. All analyses that passed the omnibus test were further assessed by Tukey’s, Holm-Sidak, or Dunn’s post hoc test analysis as noted in figure legends. In all studies, the researcher was blinded by co-investigators or by the unknown genotype of each animal.

## Results

### Repeated I/R induces prolonged hypersensitivity in mice

Our previous findings have shown that male and female mice are hypersensitive following an I/R injury to the forepaw [53,55]. These results were replicated here in that when a single I/R injury was performed, both males (p=0.033) and females (p<0.0001) demonstrate acute fore paw guarding behaviors, one day following injury **(Fig. 1)**. When a second I/R injury was performed 7 days later in mice that previously experienced a single I/R, a prolonged guarding response is observed in both males (p<0.04) and females (p<0.0001). Males returned to baseline levels by Day 8; however, females still displayed significant paw guarding at the eight-day time point. This suggests that repeated I/R injury causes prolonged pain-like behaviors in both males and females, and this repeated injury model can be used to study the transition from acute to prolonged pain-like responses.

**Figure 1:**
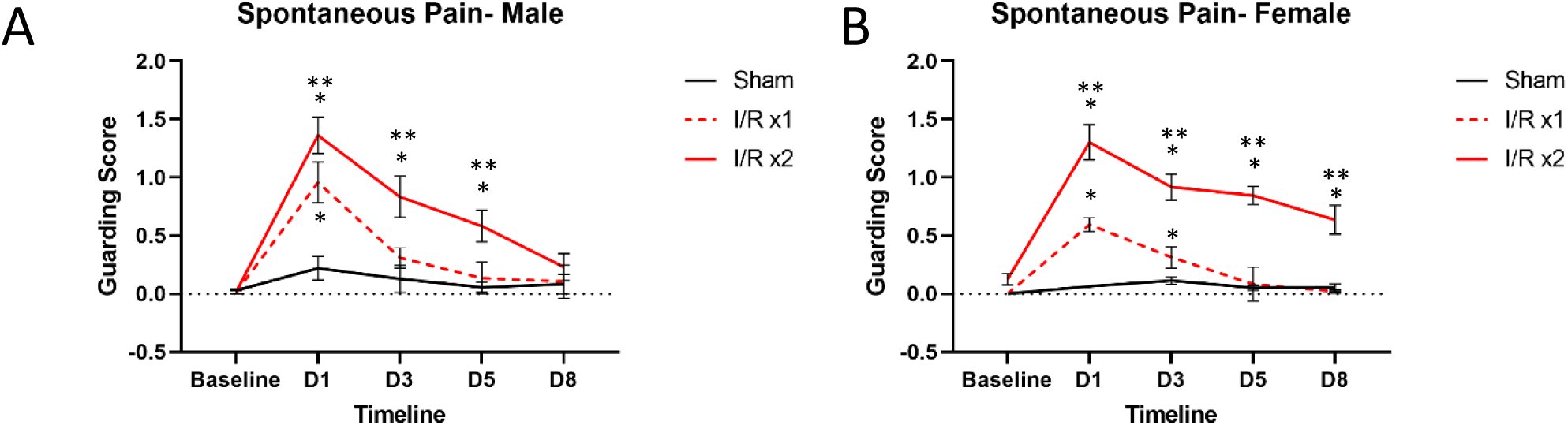
Spontaneous paw guarding behavior in mice with single or repeated I/R injury. A single I/R injury induces transient fore paw guarding for both males (A) and females (B) that lasts between 1-3d. When an I/R injury is repeated 7 days later, a prolonged phenotype is present in both males and females, but the duration is greater in females compared to males. Sham injury does not cause guarding behaviors in males or females at any time point. *p<0.05 vs baseline, **p<0.05, time-matched I/R x1 vs I/R x2, n=8 per group, 2-way RM ANOVA with Tukey’s multiple comparisons post hoc test. mean±SEM.

### Distinct muscle and DRG expression patterns are detected after I/R between sexes

Our previous findings have shown specific changes in muscle and DRG gene expression after I/R. For example, there is upregulation of muscle IL1β and its receptor, interleukin type 1 receptor 1 (IL1R1) within DRGs following a single I/R injury [54,55]. We have also reported changes in growth factors, specifically GDNF, to be upregulated in the muscle 1 day after an I/R injury as well as it’s receptor, GFRα1, increased in the affected DRGs of males [53]. We therefore wanted to determine if a similar pattern is observed after repeated I/R. The amount of GDNF and IL1β was therefore quantified in the muscle one day after the second I/R. We found that males have an upregulation of GDNF (p=0.0114) but not IL1β, while females have an upregulation of IL1β (p=0.0122) but not GDNF **(Fig. 2A)**. Immunohistochemical analysis of muscle tissue from males and females with repeated I/R confirm these findings **(Fig. 2B)**. These results corresponded to the upregulation of the GDNF co-receptor GDNF family receptor alpha 1 (GFRα1), at 1 day post single I/R as well as 1 day post double I/R injury in male mice (p<0.05). Upregulation of ASIC3 was also observed in males at both time points (p<0.05), similar to previous literature identifying a role of ASIC3 in altering muscle afferents in pain-related changes [50,54]. No changes in TRPV1 were observed after either injury in males. In female mice, there was observed upregulation of GFRα1 only one day after the second I/R injury. However, there was an upregulation in IL1R1 both 1 day after the first and second I/R injuries (p<0.05) in females. There is also an upregulation in TRPV1 in female DRGs (p<0.05) with no changes in ASIC3 **(Fig. 2C).** Together, this suggests that distinct dynamic signaling and receptor mechanisms in the periphery may underlie observed hypersensitivity in males and females after repeated I/R.

**Figure 2:**
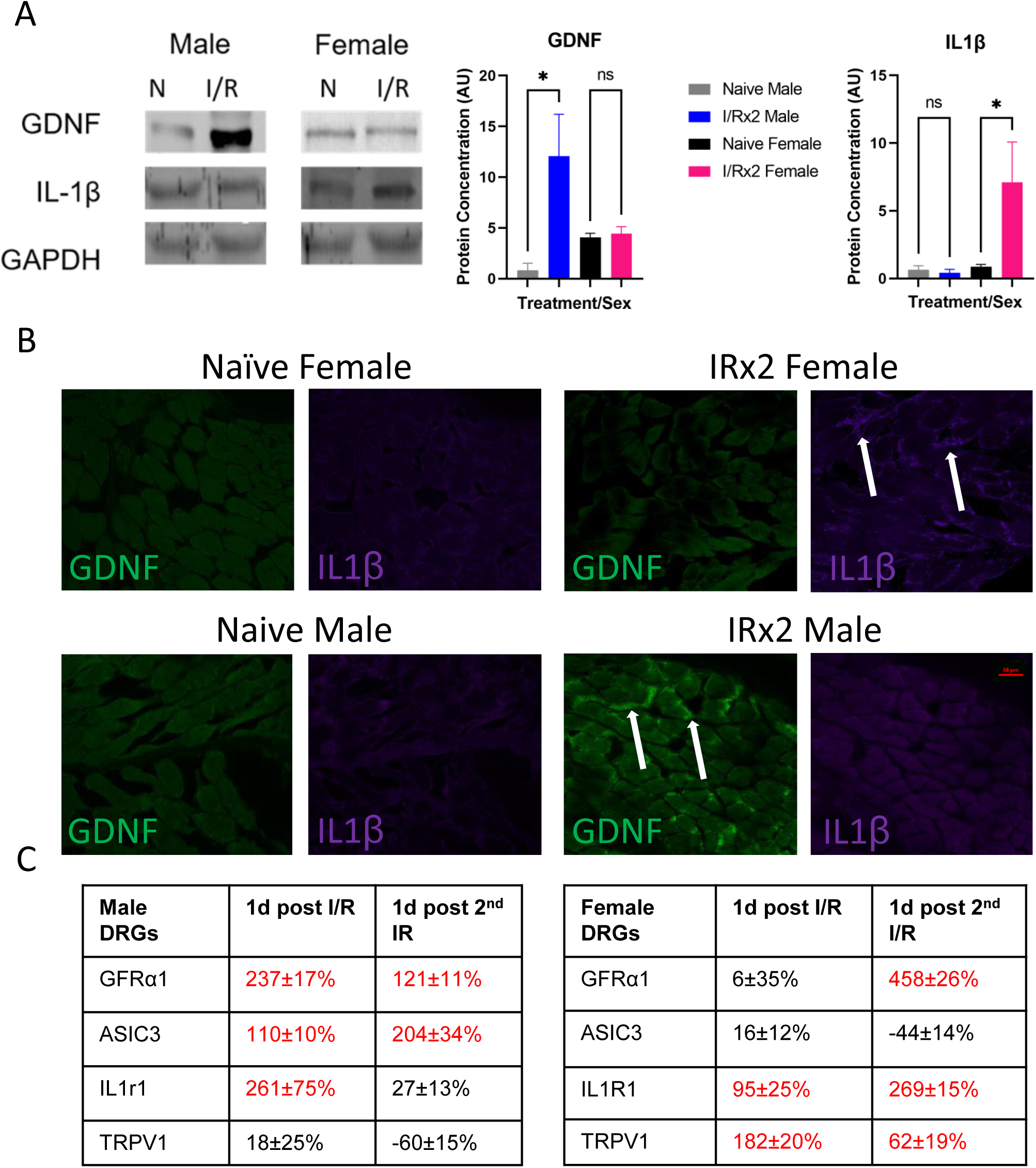
Differential muscle and DRG expression patterns in male and female mice with repeated I/R. One day following a repeated I/R injury, there is a sex-dependent upregulation of GDNF and IL1β in the muscles. Using western blotting, males display an upregulation of GDNF while females show increased IL1β following repeated I/R injury (A). Using IHC, females show increased IL1β in the muscles one day after double I/R while males display enhanced GDNF expression (B). Analysis of known receptors/channels in the DRGs in males and females following a single and double I/R injury show unique patterns of gene expression (C). A: One-way ANOVA with Holm-Sidak post-hoc; *p<0.05 between groups, n = 3-4 per group. mean±SEM. C: red indicates p<0.05 vs naïve. Values = % change vs naïve. 1-way ANOVA with Tukey’s post hoc; n=4-6 per group.

### Numerous proteins are differentially expressed between male and female DRGs

To determine possible mechanisms by which distinct receptors could be differentially altered in male and female DRGs after I/R, we used both candidate and unbiased proteomics approaches to identify potential regulatory factors in the DRGs. We found several factors that were distinctly expressed in naïve and I/R injured male and female DRGs **(Fig. 3A-D)**. Of the 1154 proteins identified and verified to be expressed in our naïve samples, 53 were differentially expressed between uninjured male and female DRGs as shown in both heat maps and volcano plots. Interestingly, of the 507 proteins detected and confirmed in our I/R injured samples, only 16 were differentially altered after I/R between sexes. We then assessed a series of RNA binding proteins in the DRGs that could also play a role in differential gene expression. We found no changes in total HuR, phosphorylated HuR or HNRNPD (aka AU-rich element RNA binding protein 1: AUF1) between male and female DRGs. However, we did find a specific increase in the phosphorylated form of AUF1 in naïve female DRGs compared to males **(Fig. 3E)**.

**Figure 3:**
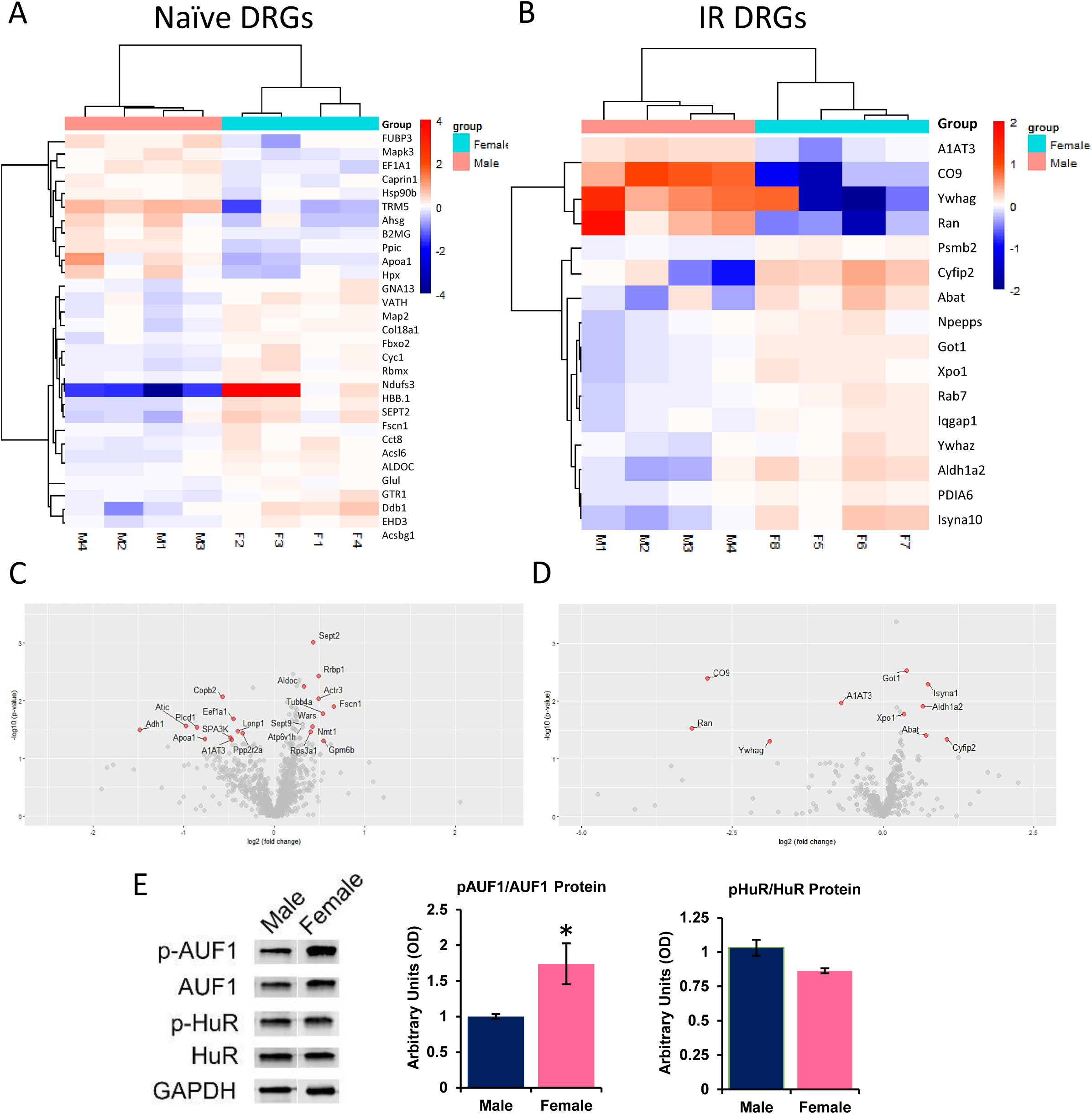
Unbiased and targeted screening analyses of protein expression in the DRGs of males and females with or without I/R injury. Numerous proteins are differentially expressed in naive (A) and I/R affected (B) male and female DRGs as assessed using comparative profiling of dorsal root ganglia tissue by SWATH nanoLC-MS/MS. 1154 total proteins were identified and verified in naïve DRGs while 507 total proteins were detected and confirmed in injured DRGs. Volcano plots comparing female/male expression of select proteins in naive (C) and I/R injured (D) male and female DRGs found to be at least 1.25x different between samples and have a p-value less than 0.05. Targeted analysis of RNA binding proteins identified phosphorylated AUF1 (p-AUF1) as significantly increased in naïve female DRGs comparted to males (E). No differences were found with HuR protein in the DRGs of males and females. One-way ANOVA; *p<0.05 vs males n=4-6 per group. mean±SEM.

### Neuronal AUF1 regulates the sex dependent development of prolonged hypersensitivity after repeated I/R injury

Since AUF1 has been linked to both stabilization and degradation of RNAs [2,10,22] and can be regulated via hormones that are known to be sex specifically expressed [4,23], we wanted to determine if these inherent differences in AUF1 activation (or expression after I/R) could underlie the differential gene expression patterns observed in male and female DRGs after injury. We therefore used our nerve-specific siRNA-mediated knock-down strategy [50,54] to inhibit the expression of AUF1 in sensory neurons after repeated I/R. We saw sex-specific effects of the AUF1 knockdown at the DRG level in that upregulation of GFRα1 and ASIC3 in the DRGs after repeated I/R in control injected male mice (PenCON+2xI/R) were not significantly affected by the nerve targeted AUF1 siRNA injections (PenAUF1+2xI/R). However, in the female DRGs, sensory neuron AUF1 knock-down prevented the repeated I/R induced upregulation of both IL1r1 and TRPV1 (p<0.05 vs naïve) after injury. Few effects were observed on GFRα1 expression in females with repeated I/R **(Figs. 4A-B)**.

**Figure 4:**
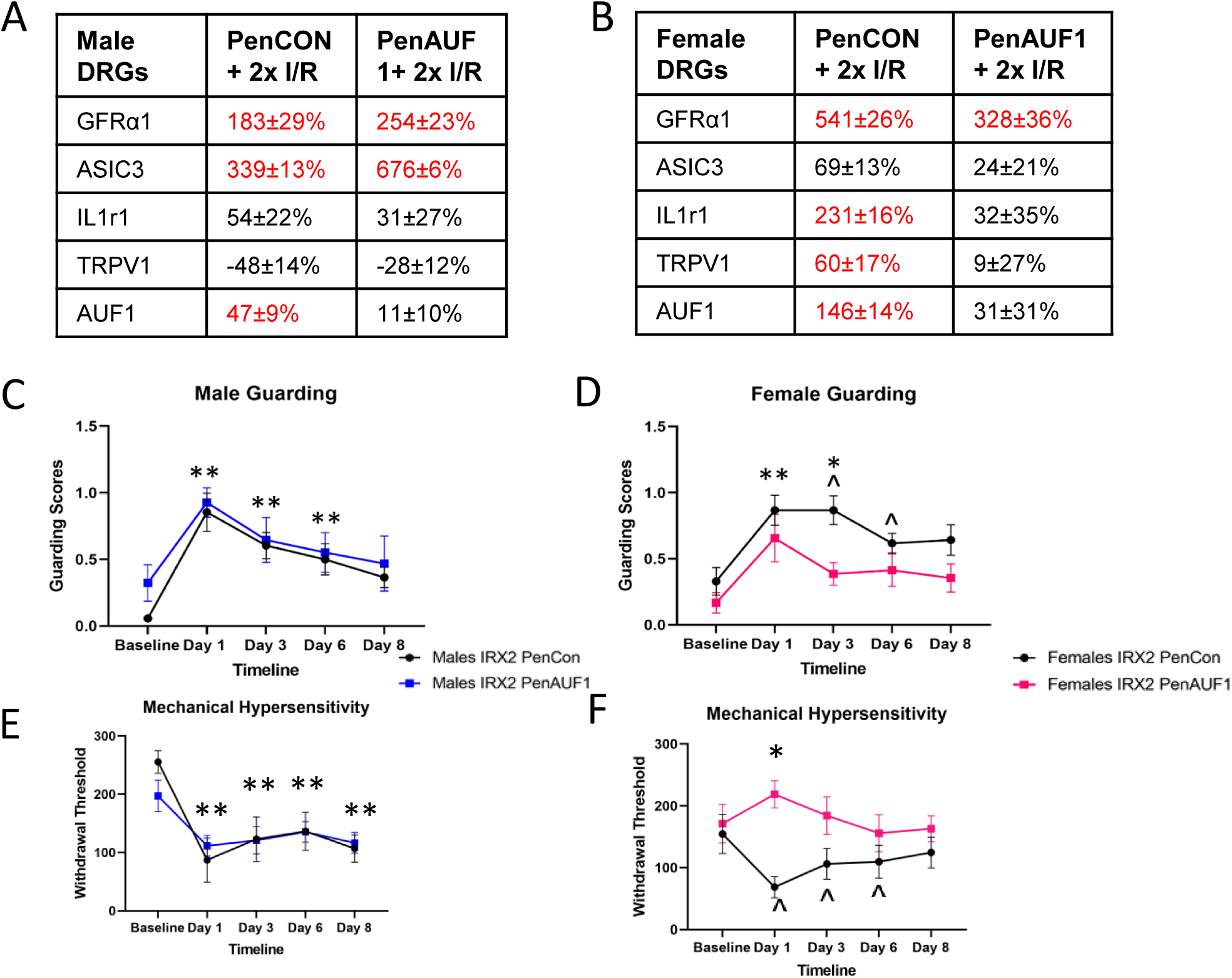
Neuronal AUF1 regualtes DRG gene expression and I/R-related pain-like behaviors in a sex dependent manner. Following nerve specific siRNA-mediated knockdown of AUF1 during repeated I/R (PenAUF1+2xI/R), no effects on select gene expression were observed in male DRGs after repeated I/R compared to controls (PenCON+2xI/R; A). However, in females, I/R-induced upregulation of IL1r1 and TRPV1 was inhibited by knockdown of AUF1 in neurons (B). No changes in I/R-induced upregulation of GFRα1 were found in either male or female DRGs with AUF1 knockdown. Although siRNA mediated knockdown of AUF1 did not alter paw guarding behaviors in males compared to controls (C), inhibition of AUF1 significanlty blunted paw guarding in females (D) following repeated I/R injury. Similar results were found regarding mechanical withdrawal thresholds in that prolonged hypersensitivity to muscle squeezing in males with dual I/R is unaffected by AUF1 inhibition (E). However, siRNA knockdown of AUF1 in females blocked prolonged mechancial hypersensitivity after repeated I/R compared to control injected mice (F). A-B: Red letters indicate p<0.05 vs naïve. Values=% change vs naïve, One-way ANOVA w/ Tukey’s post hoc, n=4-6 per group. C-F: *p<0.05 PenCon vs PenAUF1 ; **p<0.05 vs. baseline but not each other for both PenCon and PenAUF1; ^p<0.05 vs baseline for PenCon only. 2-way RM ANOVA w/ Tukey’s post hoc, n= 8-12 per group. mean±SEM.

Next, we wanted to assess if this effect extended to our behavioral measures to determine if AUF1 had an impact on hypersensitivity post repeated I/R. In both spontaneous paw guarding and mechanical hypersensitivity to fore paw muscle squeezing, control (PenCON+2xI/R) and AUF1 targeted (PenAUF1+2xI/R) males with repeated I/R were unaffected by the AUF1 knockdown **(Fig. 4C-D)**. When we analyzed female behavior, we saw significant differences between the two siRNA injected groups in that prolonged paw guarding was partially inhibited by AUF1 knockdown while mechanical hypersensitivity to muscle squeezing was completely inhibited by AUF1 targeting siRNAs in mice with repeated I/R compared to controls **(Fig. 4E-F)**. In addition, grip strength was tested in separate cohorts **(Supplementary Fig. 2)**. Normalized to weight, there was no difference in grip strength in males or females between the two groups. However, the other two behavioral measures demonstrate that AUF1 plays a significant role in female specific hypersensitivity following repeated I/R injury.

We then wanted to determine if AUF1 knockdown had any effect on TRPV1 or ASIC3 protein in the affected DRGs. Using IHC, in male mice with repeated I/R, both TRPV1 and ASIC3 were unaffected by the AUF1 knockdown **(Fig. 5)**. However, in female mice TRPV1 was significantly downregulated in the PenAUF1 group when compared to naïve and the PenCon group (p<0.0001) while ASIC3 was unaffected **(Fig. 6)**. This shows that AUF1 has the ability to regulate gene expression at the DRG level in a sex dependent manner.

**Figure 5:**
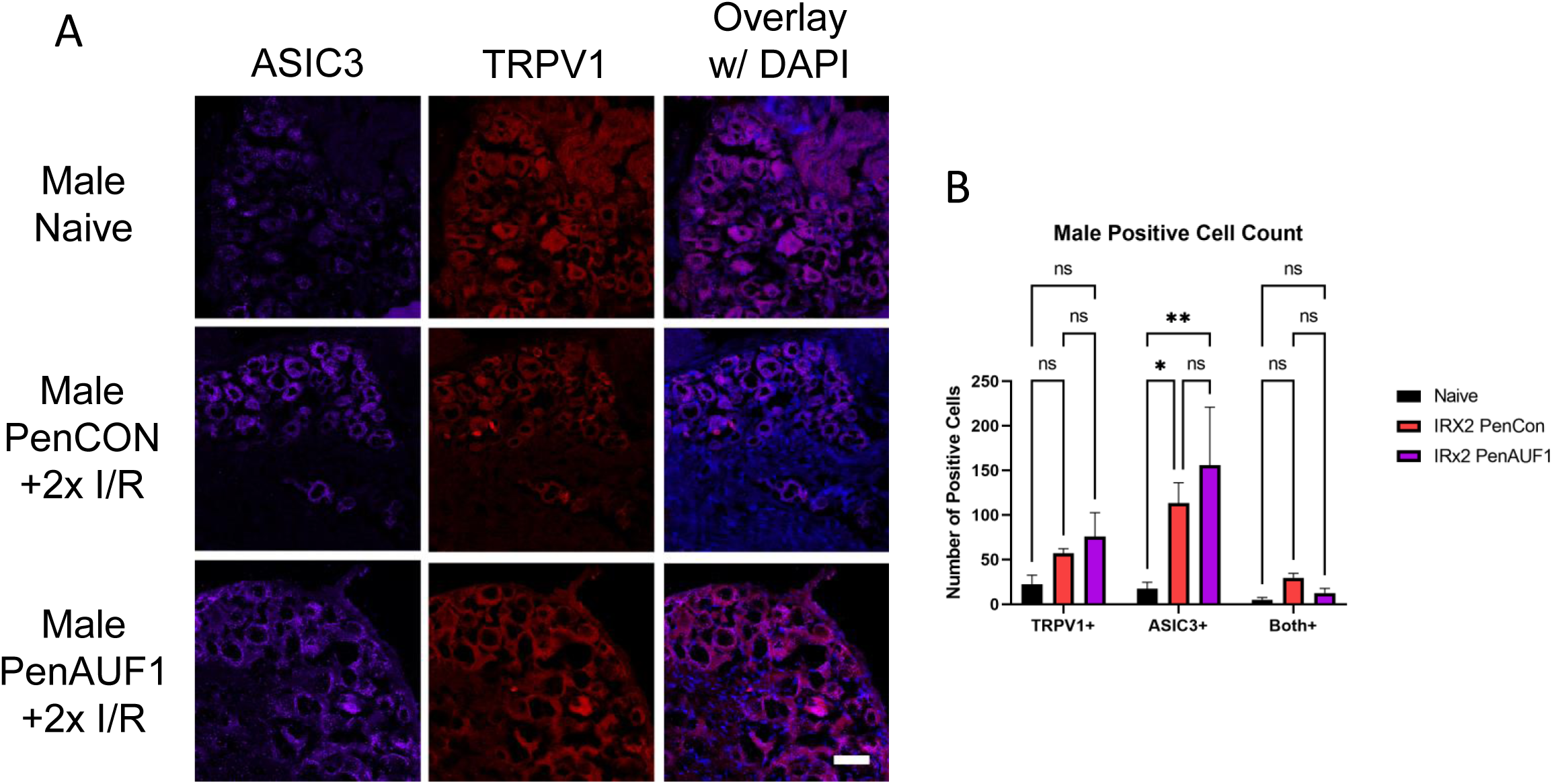
Immunohistochemical analysis of DRGs from male mice with repeated I/R and with or without AUF1 inhibition. Knockdown of AUF1 in male DRGs (A) had no effect on TRPV1 (red) or ASIC3 (purple) expression in mice with repeated I/R compared to the naïve controls (B). DAPI: blue. Scale bar, 50μm. One-way ANOVA with Tukey’s post hoc *p<0.02 between groups, n = 3-4 per group.

**Figure 6:**
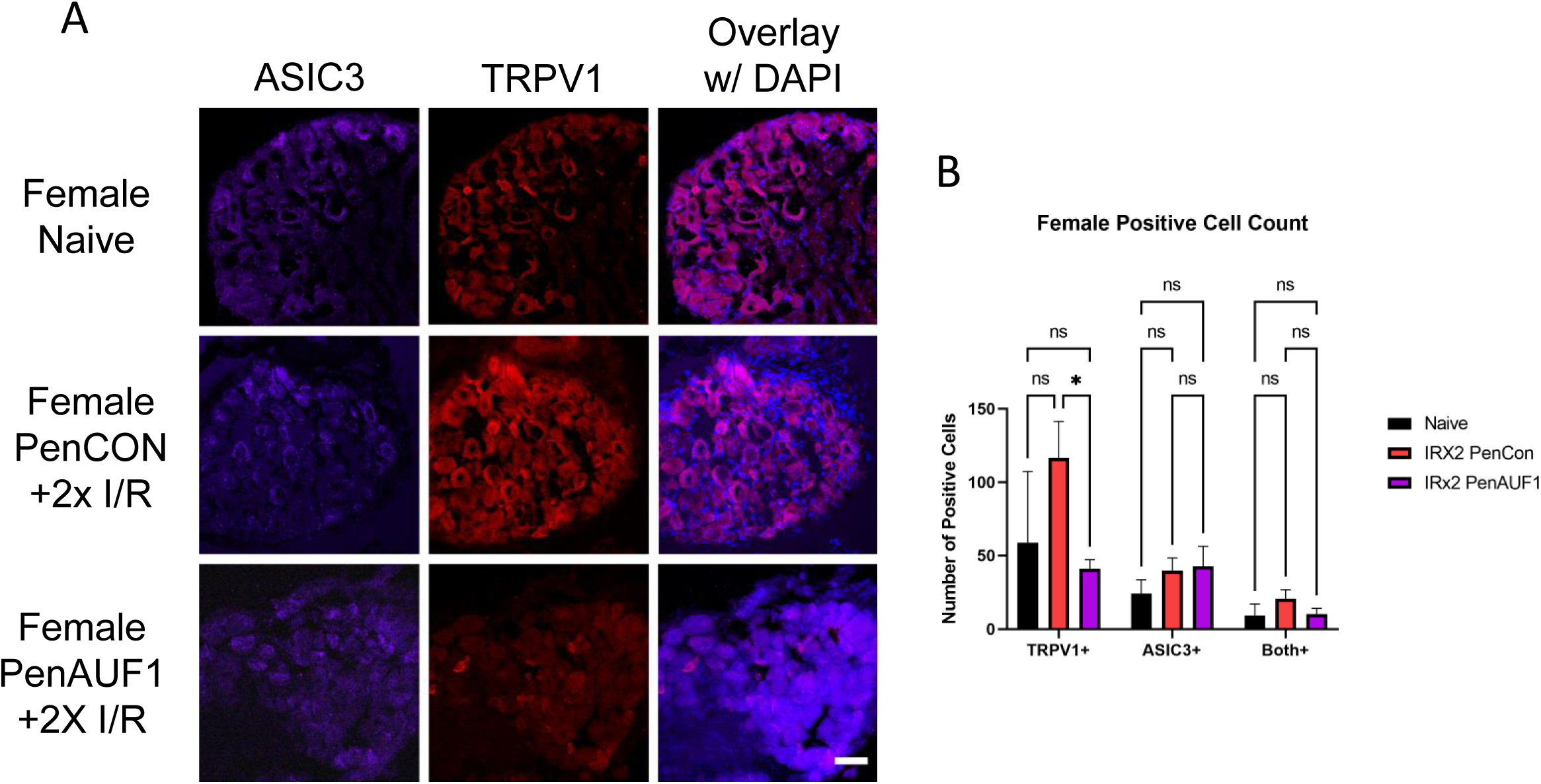
Immunohistochemical analysis of DRGs from female mice with repeated I/R and with or without AUF1 inhibition. PenCon injected female mice with repeated I/R show increased TRPV1 in the DRGs compared to naïve, which is decreased following nerve targeted, siRNA-mediated AUF1 knockdown (A,B). No effects on ASIC3 are observed in female DRGs in any condition. DAPI: blue. Scale bar, 50μm. One-way ANOVA with Tukey’s post hoc *p<0.02 between groups, n = 3-4 per group.

### Overexpression of AUF1 in males prolongs I/R related behaviors

Based on these data, we wanted to determine if upregulated AUF1 in male DRGs, would induce behavioral and gene expression changes similar to female mice with repeated I/R. Using an AAV9 containing an AUF1 overexpression construct (AUF1 OE), we injected the median and ulnar nerves in male mice and compared to control AAV9 injected males. We observed that with male paw guarding **(Fig. 7A)**, the AAV OE group had prolonged hypersensitivity compared to the AAV injected controls. Although both control and AUF1 OE injected group demonstrated prolonged hypersensitivity to mechanical stimuli after repeated I/R when compared to baseline, AUF1 OE mice did display lower baseline thresholds to muscle squeezing **(Fig. 7B)**. At the DRG level **(Fig. 7C)**, nerve specific AAV9 mediated AUF1 overexpression increased IL1r1 (195 ± 29%), TRPV1 (404 ± 53%), and GFRα1 (378 ± 14%), but seemed to have prevented ASIC3 upregulation. This demonstrates that AUF1 is playing a role in IL1r1 and TRPV1 upregulation in the DRG following repeated I/R injury and that underlying sex differences in behavioral responses to dual I/R are due in part to altered levels of phosphorylated AUF1 in sensory neurons. Overall, the data provides evidence of AUF1 modulation of hypersensitivity via possible stabilization of IL1r1 and /or TRPV1 in neurons of the DRG following repeated I/R injuries **(Fig. 8)**.

**Figure 7:**
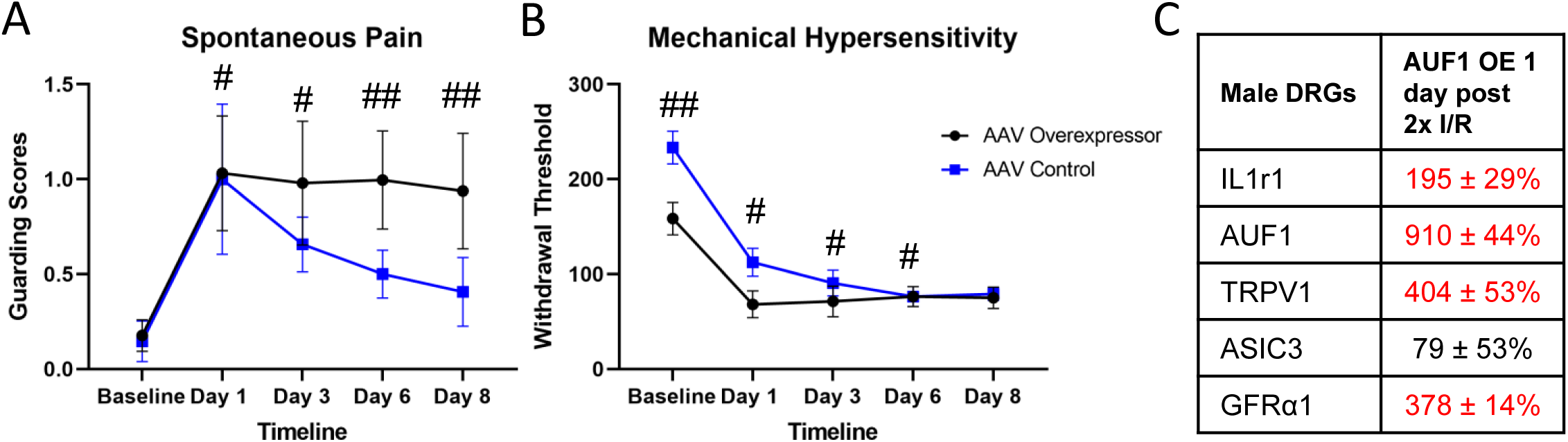
Overexpression of AUF1 in male sensory neurons prolongs I/R related pain-like behaviors and induces unique DRG gene expression. AAV mediated overexpresion of AUF1 in sensory neurons of the median and ulnar nerves enhanced spontaneous paw guarding (A) and lowered baseline mechanical withdrawal theresholds (B) after repeated I/R in males. Gene expression analysis of DRGs from males with AUF1 overexpression and repeated I/R display novel upregulation of IL1r1 and TRPV1 in addition to normally observed upregulation of GFRα1. #p<0.001 vs baseline but not each other, ##p<0.05, time-matched AAV OE vs AAV Control, 2-way RM ANOVA w/ Holm-Sidak‘s post hoc, n=8 per group mean±SEM; Red indicates p<0.05 vs AAV control in C. Values = % change vs AAV control. 1-way ANOVA, n=4 per group.

**Figure 8:**
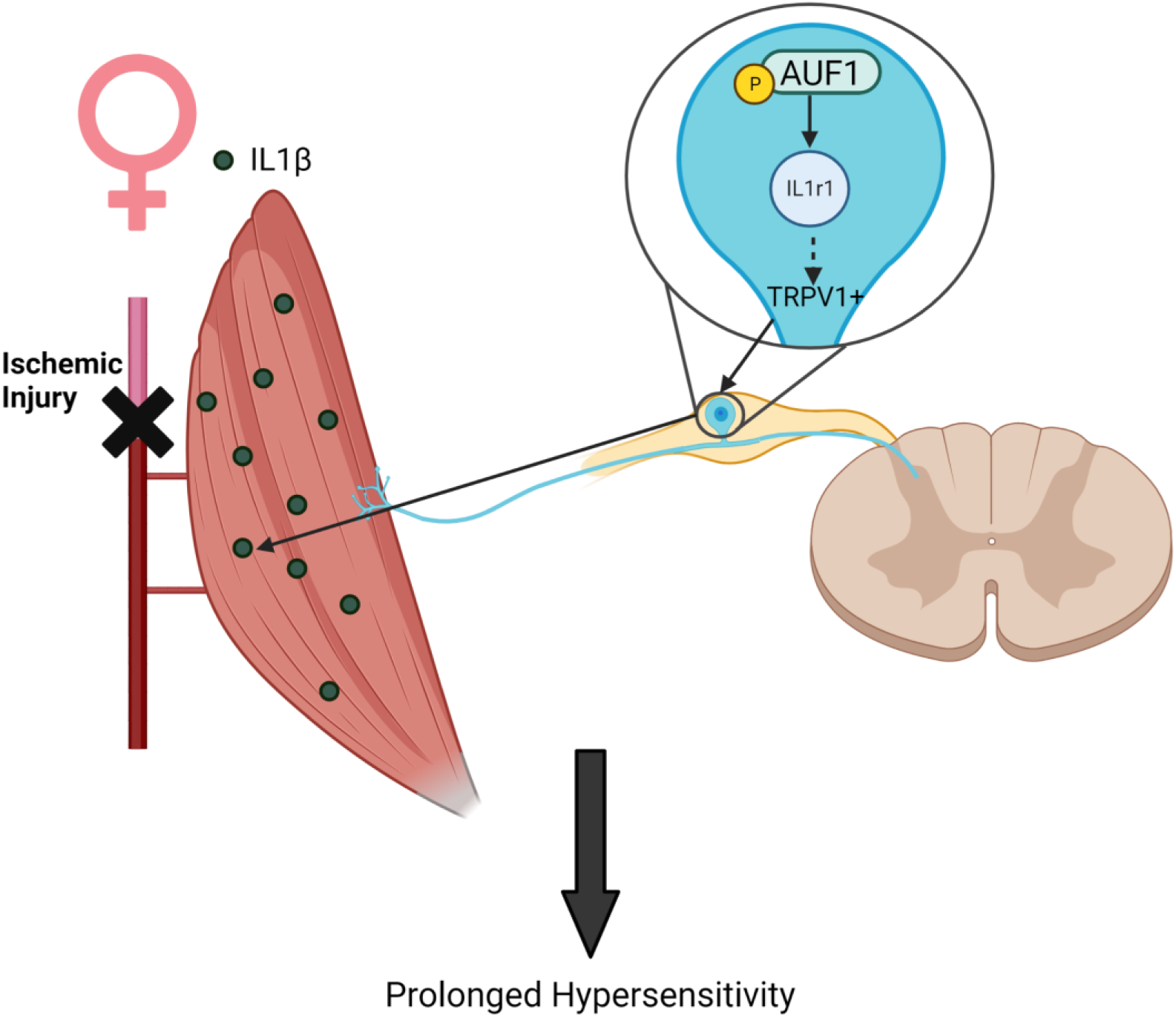
Possible mechanism by which AUF1 regulates hypersensitivity in female mice. Following repeated I/R, in females there is an increase in IL1β in the muscles with a corresponding increase in IL1r1 and TRPV1 in neurons of the DRG which may be modulated by enhanced levels of phosphorylated AUF1.

## Discussion

### Sensory neuron RNA binding proteins can modulate cytokine signaling

This study investigated the sex-specific role of sensory neuron AUF1 in a newly developed prolonged ischemic myalgia mouse model, whereby animals experience repeated I/R injuries to the forelimb. The results of our experiments found that repeated I/R induced prolonged hypersensitivity in both male and female mice [53,55], but males recovered more rapidly than females **(Fig. 1).** Our previous findings showed specific muscle and DRG gene expression changes after I/R. For example, there was upregulation of muscle IL1β and its receptor, IL1R1, within DRGs following a single I/R injury [54,55]. We also reported changes in growth factors, specifically GDNF, which was upregulated in the muscle 1 day after an I/R injury. Its receptor, GFRα1, was also increased in the affected DRGs of males [50]. We therefore wanted to determine if a similar pattern is observed after repeated I/R. Interestingly, after repeated I/R, we detected sex-specific muscle and DRG expression patterns **(Figs. 2,3,4)**. Sex-specific changes observed after I/R were similar to those found in previous reports that used other injury models including upregulation of IL1β and an increase in TRPV1+ cells [28,48,67]. In other studies from our lab, we found that ASIC3 and GFRα1 were upregulated in the DRGs of males during transient ischemic events. We previously showed that, in the muscle, IL1β and GDNF were the only growth factors and cytokines that were significantly increased following I/R injury [53]. In this study, following a repeated I/R injury, there was an upregulation of GDNF (but not IL1β) in the muscles as well as an upregulation of ASIC3 and GFRα1 in the DRG of males, similar to previous literature identifying a role of ASIC3 in altering muscle afferents in pain-related behaviors [50,54]. Females however, showed an upregulation of IL1β (but not GDNF) in the muscles and increased expression of IL1r1 and TRPV1 in the DRGs with no changes in ASIC3 **(Fig. 2)**. Together, this suggests that distinct signaling and receptor mechanisms in the periphery may underlie observed hypersensitivity in males and females after repeated I/R.

Our previous work in an acute model of ischemic injury showed that IL1β/IL1r1 signaling may have a role in regulating hypersensitivity [54]. This is partially supported by our current results. Our data showed repeated I/R increased muscle IL1β and DRG IL1r1 expression in females but not in males **(Fig. 2)**. The sex-specific activation of this pathway after repeated I/R injury may be due to a number of currently undetermined factors, but data herein suggests enhanced p-AUF1 plays a significant role **(Figs. 3,4,6)**. There is a reported connection between AUF1 and IL1β/IL1r1 signaling [9,31,58], and AUF1 has 7 binding motifs on IL1r1 mRNA. Therefore, it is possible that female specific IL1β/IL1r1 signaling in the DRGs may be mediated via differential AUF1 activation. siRNA knockdown of AUF1 in female sensory neurons during repeated I/R displayed inhibited behavioral hypersensitivity and downregulation of IL1r1 in the DRGs. There was no effect of AUF1 knockdown on male behavior or DRG gene expression however **(Figs. 4, 5).** Nevertheless, our current data showed that overexpression of AUF1 in male sensory neurons prolonged I/R-related behaviors. At the DRG level, gene expression of AUF1, IL1r1, TRPV1, and GFRα1 were upregulated in the OE mice compared to the AAV-injected controls **(Fig. 7).** Interestingly, GFRα1 was upregulated, comparable to what is seen in males following I/R injury, but injury-enhanced ASIC3 was blocked. Currently, there is no literature on interactions of AUF1 and ASIC3. We could speculate that artificially enhanced AUF1 mediated alterations in IL1R1 in male DRGs caused a compensatory inhibition of the ASIC3 upregulation, but how AUF1 overexpression in male sensory neurons modulates ASIC3 will need to be investigated in future studies. Nevertheless, these data provide further evidence of a mechanism or pathway related to IL1R1 and AUF1 in I/R-related hypersensitivity **(Fig. 8)**.

### Possible role for hormones in modulating sex specific mechanisms of ischemic myalgia development

Hormones have become a leading area of interest in sex-related DRG protein expression because of their inherent sex-specific production. Inflammatory cytokines, like IL1β and its receptor, have been reported to be involved in the pathogenesis of endometriosis, an estrogen-dependent disease often associated with chronic pain [24]. The involvement of IL1β/IL1r1 has also been shown in breast cancer studies. These studies have shown that IL1β/IL1r1 expression was mediated via estrogen [64]. Literature has also demonstrated that both estrogen and prolactin have the ability to regulate IL1β production from macrophages, while testosterone may inhibit its release [52,59].

Our model also demonstrated that females exhibited upregulation of TRPV1 gene expression as well as more TRPV1+ neurons in the DRG **(Figs. 4, 6)**. This is of particular interest as the hormone prolactin has been shown to not only directly modulate the central and peripheral neurons, but specifically TRP channels that are important to pain modulation [14, 46]. Further, it has also been shown that IL1β can regulate TRPV1 expression in sensory neurons [30,33]. Although prolactin is not the only hormone shown to be involved in pain pathways, its receptors at the neuronal level [45,46,47] seem to be a potent modulator of nociceptive processing. After repeated I/R, these hormones could play a role in the sex-specific cytokine and growth factor pathways activated in males and females. It will be important in the future to analyze various hormones in establishing sex-specific mechanisms of ischemic myalgia in animal models.

### Strengths and Limitations of the study

This report is the first to describe a role for RNA binding proteins in sensory neurons that modulate a sex specific gene expression program which regulate pain-like hypersensitivity after repeated I/R injury. A new model of prolonged hypersensitivity was also established with these studies. Using unbiased and targeted screening strategies, we uncovered a novel DRG protein with enhanced phosphorylation in females that appears to uniquely modulate IL1R1 and TRPV1 expression in DRGs. This was determined using our novel nerve-specific siRNA mediated knockdown strategy and AAV mediated overexpression technologies to reveal a role for sensory neurons specific AUF1 in these phenomena.

Although this study describes a new protein involved in ischemic myalgia-like behaviors and DRG gene expression, we were not able to determine the sensory neuron subtype that may be associated with the observed effects. The antibody used in western blotting was unable to provide adequate staining with immunocytochemical processing. It will be important in the future to determine the neurochemical and physiological phenotypes of the sensory neurons driving AUF1-mediated effects in our model of prolonged I/R-related hypersensitivity. This study also did not determine how AUF1 was sex-specifically activated in female DRGs. It is plausible that hormones could regulate this differential activation pattern [22,44,47,48,49,50] but this will need to be determined in the future. It will also be important to perform studies to determine if AUF1 or sex hormones can regulate cytokine production in other cell types that can then act on receptors expressed in sensory neurons. It has been suggested that IL1β, for example, is released from macrophages to modulate MSK pain [18,29], which we have also suggested to be important in I/R-like hypersensitivity [54]. Determining the cell types, interactions and signaling factors involved in establishing a sex specific mechanism of ischemic myalgia development will be necessary future experiments.

### Clinical Significance

Myalgia has been shown to affect multiple biological systems and their interactions [43,57,63]. Repeated I/R injury occurs in many disorders with MSK pain as an underlying feature, including peripheral vascular disease, sickle cell anemia, fibromyalgia, and complex regional pain syndrome [1,12,17,25,50,51,65]. Clinically, many MSK disorders that cause chronic pain affect men and women differently [5,36,40,48,55,60]. In many pain conditions that affect men and women equally (e.g., migraines, fibromyalgia, etc.), women exhibit greater pain severity [5,48,62,67]. This sex-specificity is seen in models of chemotherapy-induced pain [32] and in a mouse model of complex regional pain syndrome [61]. Our results extend these findings in a model of prolonged ischemic myalgia.

Preclinical studies are relatively limited in analyses of sex differences in muscle pain. The results presented here are some of the first to study sex differences in ischemic myalgias, a common cause of pain in diseases that affect women more than males [48,69]. This sex-specificity may be due to differences in gonadal hormones, neuroimmune interactions, or innate differences in noxious stimulus processing [5,8,13,39,48]. Future studies of sex differences in pain are needed to inform the development of more effective, targeted, sex-specific treatments. Current treatment strategies for diseases associated with pain and I/R injuries include opioids and exercise. However, these therapies are often not feasible due to exercise intolerance or pharmacological ineffectiveness in patients with I/R-related injuries [20,27,57]. These issues highlight the importance of better understanding how pain develops in these unique perfusion disorders. Regardless, the current report indicates a sex specific role for DRG expressed RNA binding proteins (i.e., AUF1) in I/R-related hypersensitivity.

## Acknowledgments

The authors would like to thank the University of Cincinnati Proteomics Core facility for assistance with proteomics assays. Additionally, we thank Maria Ashton, MS, RPH, MBA, Medical Writer, Department of Anesthesiology, Cincinnati Children’s Hospital Medical Center, Cincinnati, OH, for providing writing assistance, editing, and proofreading. Data will be made freely available upon request. This work was supported by grants from the NIH to MPJ (R01NS113965 and R01NS105715). The authors declare no competing financial interests.

## References

1. Aaron LA, Burke MM, Buchwald D. Overlapping conditions among patients with chronic fatigue syndrome, fibromyalgia, and temporomandibular disorder. Arch Intern Med. 2000 Jan 24;160(2):221–7. doi: 10.1001/archinte.160.2.221. PMID: 10647761.

2. Abbadi D, Yang M, Chenette DM, Andrews JJ, Schneider RJ. Muscle development and regeneration controlled by AUF1-mediated stage-specific degradation of fate-determining checkpoint mRNAs. Proc Natl Acad Sci U S A. 2019 Jun 4;116(23):11285–11290. doi: 10.1073/pnas.1901165116. Epub 2019 May 21. PMID: 31113881; PMCID: PMC6561290.

3. Ahacic K, Kåreholt I. Prevalence of musculoskeletal pain in the general Swedish population from 1968 to 2002: age, period, and cohort patterns. Pain. 2010 Oct;151(1):206–214. doi: 10.1016/j.pain.2010.07.011. Epub 2010 Aug 13. PMID: 20708842.

4. Arao Y, Kikuchi A, Kishida M, Yonekura M, Inoue A, Yasuda S, Wada S, Ikeda K, Kayama F. Stability of A+U-rich element binding factor 1 (AUF1)-binding messenger ribonucleic acid correlates with the subcellular relocalization of AUF1 in the rat uterus upon estrogen treatment. Mol Endocrinol. 2004 Sep;18(9):2255–67. doi: 10.1210/me.2004-0103. Epub 2004 Jun 10. PMID: 15192077.

5. Bartley EJ, Fillingim RB. Sex differences in pain: a brief review of clinical and experimental findings. Br J Anaesth. 2013 Jul;111(1):52–8. doi: 10.1093/bja/aet127. PMID: 23794645; PMCID: PMC3690315.

6. Blyth FM, Briggs AM, Schneider CH, Hoy DG, March LM. The Global Burden of Musculoskeletal Pain-Where to From Here? Am J Public Health. 2019 Jan;109(1):35–40. doi: 10.2105/AJPH.2018.304747. Epub 2018 Nov 29. PMID: 30495997; PMCID: PMC6301413.

7. de la Peña JB, Campbell ZT. RNA-binding proteins as targets for pain therapeutics. Neurobiol Pain. 2018 Aug-Dec;4:2-7. doi: 10.1016/j.ynpai.2018.01.003. Epub 2018 Jan 31. PMID: 30370343; PMCID: PMC6201239.

8. Cairns BE, Gazerani P. Sex-related differences in pain. Maturitas. 2009 Aug 20;63(4):292–6. doi: 10.1016/j.maturitas.2009.06.004. PMID: 19595525.

9. Carpenter S, Ricci EP, Mercier BC, Moore MJ, Fitzgerald KA. Post-transcriptional regulation of gene expression in innate immunity. Nat Rev Immunol. 2014 Jun;14(6):361–76. doi: 10.1038/nri3682. PMID: 24854588.

10. Chenette DM, Cadwallader AB, Antwine TL, Larkin LC, Wang J, Olwin BB, Schneider RJ. Targeted mRNA Decay by RNA Binding Protein AUF1 Regulates Adult Muscle Stem Cell Fate, Promoting Skeletal Muscle Integrity. Cell Rep. 2016 Aug 2;16(5):1379–1390. doi: 10.1016/j.celrep.2016.06.095. Epub 2016 Jul 21. PMID: 27452471; PMCID: PMC5323095.

11. Cimas M, Ayala A, Sanz B, Agulló-Tomás MS, Escobar A, Forjaz MJ. Chronic musculoskeletal pain in European older adults: Cross-national and gender differences. Eur J Pain. 2018 Feb;22(2):333–345. doi: 10.1002/ejp.1123. Epub 2017 Dec 12. PMID: 29235193.

12. Coderre TJ, Bennett GJ. A hypothesis for the cause of complex regional pain syndrome-type I (reflex sympathetic dystrophy): pain due to deep-tissue microvascular pathology. Pain Med. 2010 Aug;11(8):1224–38. doi: 10.1111/j.1526-4637.2010.00911.x. PMID: 20704671; PMCID: PMC4467969.

13. Dao TT, LeResche L. Gender differences in pain. J Orofac Pain. 2000 Summer;14(3):169–84; discussion 184-95. PMID: 11203754.

14. Diogenes A, Patwardhan AM, Jeske NA, Ruparel NB, Goffin V, Akopian AN, Hargreaves KM. Prolactin modulates TRPV1 in female rat trigeminal sensory neurons. J Neurosci. 2006 Aug 2;26(31):8126–36. doi: 10.1523/JNEUROSCI.0793-06.2006. PMID: 16885226; PMCID: PMC6673790.

15. Elsamadicy AA, Yang S, Sergesketter AR, Ashraf B, Charalambous L, Kemeny H, Ejikeme T, Ren X, Pagadala P, Parente B, Xie J, Lad SP. Prevalence and Cost Analysis of Complex Regional Pain Syndrome (CRPS): A Role for Neuromodulation. Neuromodulation. 2018 Jul;21(5):423–430. doi: 10.1111/ner.12691. Epub 2017 Sep 29. PMID: 28961359; PMCID: PMC5876058.

16. El-Tallawy SN, Nalamasu R, Salem GI, LeQuang JAK, Pergolizzi JV, Christo PJ. Management of Musculoskeletal Pain: An Update with Emphasis on Chronic Musculoskeletal Pain. Pain Ther. 2021 Jun;10(1):181–209. doi: 10.1007/s40122-021-00235-2. Epub 2021 Feb 11. PMID: 33575952; PMCID: PMC8119532.

17. Elvin A, Siösteen AK, Nilsson A, Kosek E. Decreased muscle blood flow in fibromyalgia patients during standardised muscle exercise: a contrast media enhanced colour Doppler study. Eur J Pain. 2006 Feb;10(2):137–44. doi: 10.1016/j.ejpain.2005.02.001. PMID: 16310717.

18. Ernberg, M., Christidis, N., Ghafouri, B., Bileviciute-Ljungar, I., Lofgren, M., Larsson, A., Palstam, A., Bjersing, J., Mannerkorpi, K., Kosek, E., Gerdle, B., 2016. Effects of 15 weeks of resistance exercise on pro-inflammatory cytokine levels in the vastus lateralis muscle of patients with fibromyalgia. Arthritis Res. Ther. 18 (1), 137.

19. Fillingim RB, King CD, Ribeiro-Dasilva MC, Rahim-Williams B, Riley JL 3rd. Sex, gender, and pain: a review of recent clinical and experimental findings. J Pain. 2009 May;10(5):447–85. doi: 10.1016/j.jpain.2008.12.001. PMID: 19411059; PMCID: PMC2677686.

20. Hauet T, Pisani DF. New Strategies Protecting from Ischemia/Reperfusion. Int J Mol Sci. 2022 Dec 14;23(24):15867. doi: 10.3390/ijms232415867. PMID: 36555508; PMCID: PMC9779635.

21. Hawker GA. The assessment of musculoskeletal pain. Clin Exp Rheumatol. 2017 Sep-Oct;35 Suppl 107(5):8-12. Epub 2017 Sep 28. PMID: 28967361.

22. Hendrayani SF, Al-Khalaf HH, Aboussekhra A. The cytokine IL-6 reactivates breast stromal fibroblasts through transcription factor STAT3-dependent up-regulation of the RNA-binding protein AUF1. J Biol Chem. 2014 Nov 7;289(45):30962–76. doi: 10.1074/jbc.M114.594044. Epub 2014 Sep 17. PMID: 25231991; PMCID: PMC4223303.

23. Ing NH, Massuto DA, Jaeger LA. Estradiol up-regulates AUF1p45 binding to stabilizing regions within the 3’-untranslated region of estrogen receptor alpha mRNA. J Biol Chem. 2008 Jan 18;283(3):1764–1772. doi: 10.1074/jbc.M704745200. Epub 2007 Nov 19. PMID: 18029355.

24. Kato T, Yasuda K, Matsushita K, Ishii KJ, Hirota S, Yoshimoto T, Shibahara H. Interleukin-1/-33 Signaling Pathways as Therapeutic Targets for Endometriosis. Front Immunol. 2019 Aug 22;10:2021. doi: 10.3389/fimmu.2019.02021. PMID: 31507610; PMCID: PMC6714064.

25. Katz DL, Greene L, Ali A, Faridi Z. The pain of fibromyalgia syndrome is due to muscle hypoperfusion induced by regional vasomotor dysregulation. Med Hypotheses. 2007;69(3):517–25. doi: 10.1016/j.mehy.2005.10.037. Epub 2007 Mar 21. PMID: 17376601.

26. Khoutorsky A, Bonin RP, Sorge RE, Gkogkas CG, Pawlowski SA, Jafarnejad SM, Pitcher MH, Alain T, Perez-Sanchez J, Salter EW, Martin L, Ribeiro-da-Silva A, De Koninck Y, Cervero F, Mogil JS, Sonenberg N. Translational control of nociception via 4E-binding protein 1. Elife. 2015 Dec 18;4:e12002. doi: 10.7554/eLife.12002. PMID: 26678009; PMCID: PMC4695384.

27. Kindler LL, Valencia C, Fillingim RB, George SZ. Sex differences in experimental and clinical pain sensitivity for patients with shoulder pain. Eur J Pain. 2011 Feb;15(2):118–23. doi: 10.1016/j.ejpain.2010.06.001. Epub 2010 Jul 2. PMID: 20598598; PMCID: PMC2965801.

28. Lenert ME, Avona A, Garner KM, Barron LR, Burton MD. Sensory Neurons, Neuroimmunity, and Pain Modulation by Sex Hormones. Endocrinology. 2021 Aug 1;162(8):bqab109. doi: 10.1210/endocr/bqab109. PMID: 34049389; PMCID: PMC8237991.

29. Lesnak JB, Berardi G, Sluka KA. Influence of routine exercise on the peripheral immune system to prevent and alleviate pain. Neurobiol Pain. 2023 Mar 21;13:100126. doi: 10.1016/j.ynpai.2023.100126. PMID: 37179769; PMCID: PMC10173010.

30. Liu L, Yang TM, Liedtke W, Simon SA. Chronic IL-1beta signaling potentiates voltage-dependent sodium currents in trigeminal nociceptive neurons. J Neurophysiol. 2006 Mar;95(3):1478–90. doi: 10.1152/jn.00509.2005. Epub 2005 Nov 30. PMID: 16319216.

31. Lu JY, Sadri N, Schneider RJ. Endotoxic shock in AUF1 knockout mice mediated by failure to degrade proinflammatory cytokine mRNAs. Genes Dev. 2006 Nov 15;20(22):3174–84. doi: 10.1101/gad.1467606. Epub 2006 Nov 3. PMID: 17085481; PMCID: PMC1635151.

32. Luo X, Chen O, Wang Z, Bang S, Ji J, Lee SH, Huh Y, Furutani K, He Q, Tao X, Ko MC, Bortsov A, Donnelly CR, Chen Y, Nackley A, Berta T, Ji RR. IL-23/IL-17A/TRPV1 axis produces mechanical pain via macrophage-sensory neuron crosstalk in female mice. Neuron. 2021 Sep 1;109(17):2691–2706.e5. doi: 10.1016/j.neuron.2021.06.015. Epub 2021 Jul 19. PMID: 34473953; PMCID: PMC8425601.

33. Mailhot B, Christin M, Tessandier N, Sotoudeh C, Bretheau F, Turmel R, Pellerin È, Wang F, Bories C, Joly-Beauparlant C, De Koninck Y, Droit A, Cicchetti F, Scherrer G, Boilard E, Sharif-Naeini R, Lacroix S. Neuronal interleukin-1 receptors mediate pain in chronic inflammatory diseases. J Exp Med. 2020 Sep 7;217(9):e20191430. doi: 10.1084/jem.20191430. PMID: 32573694; PMCID: PMC7478735.

34. Mapplebeck JCS, Beggs S, Salter MW. Sex differences in pain: a tale of two immune cells. Pain. 2016 Feb;157 Suppl 1:S2–S6. doi: 10.1097/j.pain.0000000000000389. PMID: 26785152.

35. Meints SM, Cortes A, Morais CA, Edwards RR. Racial and ethnic differences in the experience and treatment of noncancer pain. Pain Manag. 2019 May;9(3):317–334. doi: 10.2217/pmt-2018-0030. Epub 2019 May 29. PMID: 31140916; PMCID: PMC6587104.12

36. Melchior M, Poisbeau P, Gaumond I, Marchand S. Insights into the mechanisms and the emergence of sex-differences in pain. Neuroscience. 2016 Dec 3;338:63–80. doi: 10.1016/j.neuroscience.2016.05.007. Epub 2016 May 12. PMID: 27180284.

37. Mense S. Muscle pain: mechanisms and clinical significance. Dtsch Arztebl Int. 2008 Mar;105(12):214–9. doi: 10.3238/artzebl.2008.0214. Epub 2008 Mar 21. PMID: 19629211; PMCID: PMC2696782.

38. Meucci RD, Fassa AG, Faria NM. Prevalence of chronic low back pain: systematic review. Rev Saude Publica. 2015;49:1. doi: 10.1590/S0034-8910.2015049005874. Epub 2015 Oct 20. PMID: 26487293; PMCID: PMC4603263.

39. Mogil JS. Sex differences in pain and pain inhibition: multiple explanations of a controversial phenomenon. Nat Rev Neurosci. 2012 Dec;13(12):859–66. doi: 10.1038/nrn3360. PMID: 23165262.

40. Mogil JS, Bailey AL. Sex and gender differences in pain and analgesia. Prog Brain Res. 2010;186:141–57. doi: 10.1016/B978-0-444-53630-3.00009-9. PMID: 21094890.

41. Meints SM, Cortes A, Morais CA, Edwards RR. Racial and ethnic differences in the experience and treatment of noncancer pain. Pain Manag. 2019 May;9(3):317–334. doi: 10.2217/pmt-2018-0030. Epub 2019 May 29. PMID: 31140916; PMCID: PMC6587104.

42. Morgan CP, Bale TL. Sex differences in microRNA regulation of gene expression: no smoke, just miRs. Biol Sex Differ. 2012 Sep 26;3(1):22. doi: 10.1186/2042-6410-3-22. PMID: 23009289; PMCID: PMC3507674.

43. Murphy MN, Mizuno M, Mitchell JH, Smith SA. Cardiovascular regulation by skeletal muscle reflexes in health and disease. Am J Physiol Heart Circ Physiol. 2011 Oct;301(4):H1191-204. doi: 10.1152/ajpheart.00208.2011. Epub 2011 Aug 12. PMID: 21841019; PMCID: PMC3197431.

44. Nagaoka K, Tanaka T, Imakawa K, Sakai S. Involvement of RNA binding proteins AUF1 in mammary gland differentiation. Exp Cell Res. 2007 Aug 1;313(13):2937–45. doi: 10.1016/j.yexcr.2007.04.017. Epub 2007 Apr 20. PMID: 17512931.

45. Paige C, Barba-Escobedo PA, Mecklenburg J, Patil M, Goffin V, Grattan DR, Dussor G, Akopian AN, Price TJ. Neuroendocrine Mechanisms Governing Sex Differences in Hyperalgesic Priming Involve Prolactin Receptor Sensory Neuron Signaling. J Neurosci. 2020 Sep 9;40(37):7080–7090. doi: 10.1523/JNEUROSCI.1499-20.2020. Epub 2020 Aug 12. PMID: 32801151; PMCID: PMC7480243.

46. Patil MJ, Ruparel SB, Henry MA, Akopian AN. Prolactin regulates TRPV1, TRPA1, and TRPM8 in sensory neurons in a sex-dependent manner: Contribution of prolactin receptor to inflammatory pain. Am J Physiol Endocrinol Metab. 2013 Nov 1;305(9):E1154–64. doi: 10.1152/ajpendo.00187.2013. Epub 2013 Sep 10. PMID: 24022869; PMCID: PMC3840203.

47. Patil M, Belugin S, Mecklenburg J, Wangzhou A, Paige C, Barba-Escobedo PA, Boyd JT, Goffin V, Grattan D, Boehm U, Dussor G, Price TJ, Akopian AN. Prolactin Regulates Pain Responses via a Female-Selective Nociceptor-Specific Mechanism. iScience. 2019 Oct 25;20:449–465. doi: 10.1016/j.isci.2019.09.039. Epub 2019 Oct 1. PMID: 31627131; PMCID: PMC6818331.

48. Queme LF, Jankowski MP. Sex differences and mechanisms of muscle pain. Curr Opin Physiol. 2019 Oct;11:1–6. doi: 10.1016/j.cophys.2019.03.006. Epub 2019 Apr 2. PMID: 31245656; PMCID: PMC6594402.

49. Queme LF, Ross JL, Lu P, Hudgins RC, Jankowski MP. Dual Modulation of Nociception and Cardiovascular Reflexes during Peripheral Ischemia through P2Y1 Receptor-Dependent Sensitization of Muscle Afferents. J Neurosci. 2016 Jan 6;36(1):19–30. doi: 10.1523/JNEUROSCI.2856-15.2016. PMID: 26740646; PMCID: PMC4701959.

50. Queme LF, Weyler AA, Cohen ER, Hudgins RC, Jankowski MP. A dual role for peripheral GDNF signaling in nociception and cardiovascular reflexes in the mouse. Proc Natl Acad Sci U S A. 2020 Jan 7;117(1):698–707. doi: 10.1073/pnas.1910905116. Epub 2019 Dec 17. PMID: 31848242; PMCID: PMC6955322.

51. Queme LF, Ross JL, Jankowski MP. Peripheral Mechanisms of Ischemic Myalgia. Front Cell Neurosci. 2017 Dec 22;11:419. doi: 10.3389/fncel.2017.00419. PMID: 29311839; PMCID: PMC5743676.

52. Rosen S, Ham B, Mogil JS. Sex differences in neuroimmunity and pain. J Neurosci Res. 2017 Jan 2;95(1-2):500–508. doi: 10.1002/jnr.23831. PMID: 27870397.

53. Ross JL, Queme LF, Shank AT, Hudgins RC, Jankowski MP. Sensitization of group III and IV muscle afferents in the mouse after ischemia and reperfusion injury. J Pain. 2014 Dec;15(12):1257–70. doi: 10.1016/j.jpain.2014.09.003. PMID: 25245401; PMCID: PMC4302035.

54. Ross JL, Queme LF, Cohen ER, Green KJ, Lu P, Shank AT, An S, Hudgins RC, Jankowski MP. Muscle IL1β Drives Ischemic Myalgia via ASIC3-Mediated Sensory Neuron Sensitization. J Neurosci. 2016 Jun 29;36(26):6857–71. doi: 10.1523/JNEUROSCI.4582-15.2016. PMID: 27358445; PMCID: PMC4926236.

55. Ross JL, Queme LF, Lamb JE, Green KJ, Jankowski MP. Sex differences in primary muscle afferent sensitization following ischemia and reperfusion injury. Biol Sex Differ. 2018 Jan 3;9(1):2. doi: 10.1186/s13293-017-0163-5. PMID: 29298725; PMCID: PMC5751812.

56. Sarkar S, Sinsimer KS, Foster RL, Brewer G, Pestka S. AUF1 isoform-specific regulation of anti-inflammatory IL10 expression in monocytes. J Interferon Cytokine Res. 2008 Nov;28(11):679–91. doi: 10.1089/jir.2008.0028. PMID: 18844578; PMCID: PMC2956575.

57. Siracusa R, Paola RD, Cuzzocrea S, Impellizzeri D. Fibromyalgia: Pathogenesis, Mechanisms, Diagnosis and Treatment Options Update. Int J Mol Sci. 2021 Apr 9;22(8):3891. doi: 10.3390/ijms22083891. PMID: 33918736; PMCID: PMC8068842.

58. Sirenko OI, Lofquist AK, DeMaria CT, Morris JS, Brewer G, Haskill JS. Adhesion-dependent regulation of an A+U-rich element-binding activity associated with AUF1. Mol Cell Biol. 1997 Jul;17(7):3898–906. doi: 10.1128/MCB.17.7.3898. PMID: 9199324; PMCID: PMC232242.

59. Smith JA, Das A, Butler JT, Ray SK, Banik NL. Estrogen or estrogen receptor agonist inhibits lipopolysaccharide induced microglial activation and death. Neurochem Res. 2011 Sep;36(9):1587–93. doi: 10.1007/s11064-010-0336-7. Epub 2010 Dec 3. Erratum in: Neurochem Res. 2011 Sep;36(9):1715. PMID: 21127968; PMCID: PMC3951894.

60. Sorge RE, Totsch SK. Sex Differences in Pain. J Neurosci Res. 2017 Jun;95(6):1271–1281. doi: 10.1002/jnr.23841. Epub 2016 Jul 25. PMID: 27452349.

61. Tajerian M, Sahbaie P, Sun Y, Leu D, Yang HY, Li W, Huang TT, Kingery W, David Clark J. Sex differences in a Murine Model of Complex Regional Pain Syndrome. Neurobiol Learn Mem. 2015 Sep;123:100–9. doi: 10.1016/j.nlm.2015.06.004. Epub 2015 Jun 9. PMID: 26070658; PMCID: PMC4530062.

62. Traub RJ, Ji Y. Sex differences and hormonal modulation of deep tissue pain. Front Neuroendocrinol. 2013 Oct;34(4):350–66. doi: 10.1016/j.yfrne.2013.07.002. Epub 2013 Jul 17. PMID: 23872333; PMCID: PMC3830473.

63. Trouvin AP, Perrot S. New concepts of pain. Best Pract Res Clin Rheumatol. 2019 Jun;33(3):101415. doi: 10.1016/j.berh.2019.04.007. Epub 2019 May 13. PMID: 31703792.

64. Al-Tweigeri T, AlRaouji NN, Tulbah A, Arafah M, Aboussekhra M, Al-Mohanna F, Gad AM, Eldali AM, Elhassan TA, Aboussekhra A. High AUF1 level in stromal fibroblasts promotes carcinogenesis and chemoresistance and predicts unfavorable prognosis among locally advanced breast cancer patients. Breast Cancer Res. 2022 Jul 11;24(1):46. doi: 10.1186/s13058-022-01543-x. PMID: 35821051; PMCID: PMC9275022.

65. Vierck CJ. A mechanism-based approach to prevention of and therapy for fibromyalgia. Pain Res Treat. 2012;2012:951354. doi: 10.1155/2012/951354. Epub 2011 Oct 2. PMID: 22110947; PMCID: PMC3200141.

66. White EJ, Brewer G, Wilson GM. Post-transcriptional control of gene expression by AUF1: mechanisms, physiological targets, and regulation. Biochim Biophys Acta. 2013 Jun-Jul;1829(6-7):680-8. doi: 10.1016/j.bbagrm.2012.12.002. Epub 2012 Dec 14. PMID: 23246978; PMCID: PMC3664190.

67. Wiesenfeld-Hallin Z. Sex differences in pain perception. Gend Med. 2005 Sep;2(3):137–45. doi: 10.1016/s1550-8579(05)80042-7. PMID: 16290886.

68. Wijnhoven HA, de Vet HC, Picavet HS. Explaining sex differences in chronic musculoskeletal pain in a general population. Pain. 2006 Sep;124(1-2):158–66. doi: 10.1016/j.pain.2006.04.012. Epub 2006 May 22. PMID: 16716517.

69. Wolfe F, Walitt B, Perrot S, Rasker JJ, Häuser W. Fibromyalgia diagnosis and biased assessment: Sex, prevalence and bias. PLoS One. 2018 Sep 13;13(9):e0203755. doi: 10.1371/journal.pone.0203755. PMID: 30212526; PMCID: PMC6136749.

70. Zucconi BE, Wilson GM. Modulation of neoplastic gene regulatory pathways by the RNA-binding factor AUF1. Front Biosci (Landmark Ed). 2011 Jun 1;16(6):2307–25. doi: 10.2741/3855. PMID: 21622178; PMCID: PMC3589912.

