## Supplementary Figures for "Sex specific role of RNA-binding protein, AUF1, on prolonged hypersensitivity after repetitive ischemia with reperfusion injury"

Supplementary Figure 1

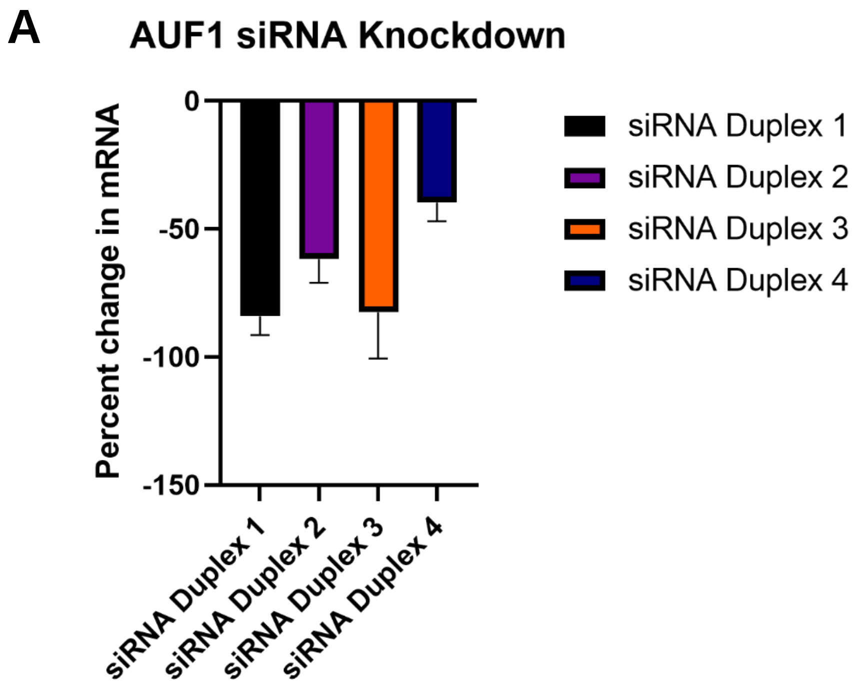

Supplementary Figure 2

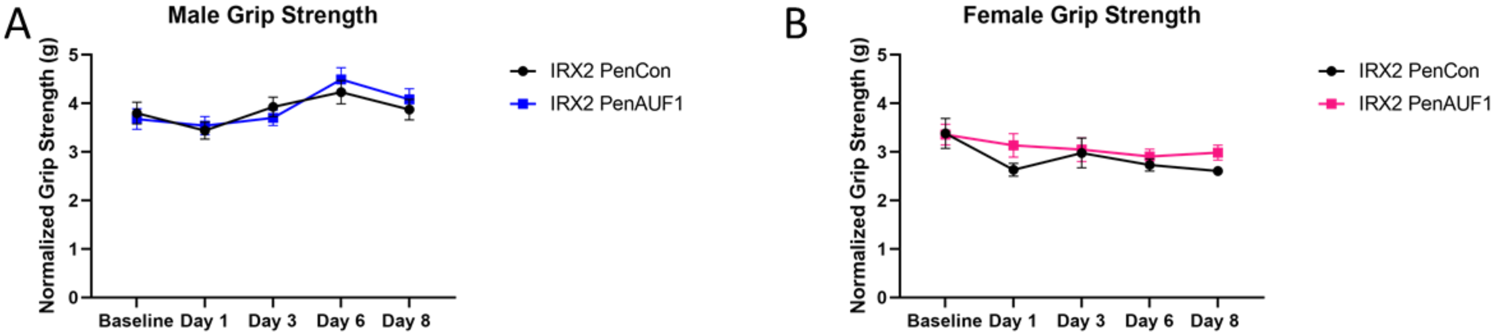

### Supplementary Table 1

| Gene | Forward | Reverse |
|---|---|---|
| GAPDH | ATGTGTCCGTCGTGGATCTGA | ATGCCTGCTTCACCACCTTCTT |
| GFRα1 | GTGTGCAGATGCTGTGGACTAG | TTCAGTGCTTCACACGCACTTG |
| ASIC3 | CTCAACTATGAGGCCGTGGAACAA | TAAGCAGGCTGGCTCCGATAAACA |
| IL1r1 | AGGAATGTGGCTGAAGAGCACAGA | ACTCGTGTGACCGGATATTGCTTC |
| TRPV1 | TTCCTGCAGAAGAGCAAGAAGC | CCCATTGTGCAGATTGAGCAT |
| AUF1 | AGATCGACGCCAGTAAGA | CAGCTAAGGCCTCCTATAAAC |
