## Supplementary Figure Legends for "Sex specific role of RNA-binding protein, AUF1, on prolonged hypersensitivity after repetitive ischemia with reperfusion injury"

**Supplementary Figure 1: Realtime PCR analysis of siRNA mediated knockdown efficiency of AUF1 in Neuro2A cells.** Using 4 distinct duplexes targeting AUF1, it was determined that duplex #1 had the most efficient knockdown capacity. This was used for conjugation to Penetratin-1 and used in remaining studies *in vivo*.

**Supplementary Figure 2: AUF1 knockdown does not alter grip strength in males or females.** Following AUF1 knockdown, no differences in grip strength are found in male (A) or female mice (B) with repeated I/R. 2-way RM ANOVA w/ Holm-Sidak’s post hoc, n=4 per group, p>0.05.

**Supplementary Table: Primer information for genes tested using realtime PCR.**
